# Flagellar Motility During E. Coli Biofilm Formation Provides a Competitive Disadvantage Which Recedes in the Presence of Co-Colonizers

**DOI:** 10.1101/2022.04.05.487170

**Authors:** Wafa Ben Youssef, Maxime Deforet, Amaury Monmeyran, Nelly Henry

## Abstract

In nature, bacteria form biofilms in very diverse environments, involving a range of specific properties and exhibiting competitive advantages for surface colonization. However, the underlying mechanisms are difficult to decipher. In particular, the contribution of cell flagellar motility to biofilm formation remains unclear. Here, we examined the ability of motile and nonmotile *E. coli* cells to form a biofilm in a well-controlled geometry, both in a simple situation involving a single-species biofilm and in the presence of co-colonizers. Using a millifluidic channel, we determined that motile cells have a clear disadvantage in forming a biofilm, exhibiting a long delay as compared to nonmotile cells. By monitoring biofilm development in real time, we observed that the decisive impact of flagellar motility on biofilm formation consists in the alteration of surface access time. Implementing a simple mathematical model to calculate surface access velocity for both motile (diffusion) and nonmotile (settling) cells, we discovered that the competitive advantage strongly depends on the geometry of the environment to be colonized. Interestingly, this advantage reverts for the smallest geometries, with a crossover at a height of 100 μm. We also report that the difference between and nonmotile cells in the ability to form a biofilm diminishes in the presence of co-colonizers, which could be due to motility inhibition through the consumption of key resources by the co-colonizers. We conclude that the impact of flagellar motility on surface colonization closely depends on the environment geometry and the population features, suggesting a unifying vision of the role of cell motility in surface colonization and biofilm formation.

## Introduction

Biofilms represent the preferred lifestyle for bacteria^1^. In these three-dimensional structures, the aggregated cells prosper in a self-produced polymer extracellular matrix that protects them from shear stress, grazers and biocides^2,3^. The formation of a biofilm is a highly multifactorial process in which the cell properties and the details of the environment are both important, bringing about a tremendous diversity in behavior.

In the planktonic state, most bacteria swim in a series of runs and tumbles and rotate their flagella assembled in bundles^4–6^, which provides a significant adaptive advantage in nutrient search^7^. The importance of this cell motility to biofilm formation has been investigated in a large spectrum of bacterial species and environmental conditions, resulting in differing viewpoints. For example, early research determined that motility was crucial for biofilm development in *Pseudomonas aeruginosa^8^, Listeria monocytogenes^9^* and *Escherichia coli^10,11^.* Furthermore, flagellar motility is often associated with increased virulence in pathogenic species, with motile bacteria exhibiting facilitated host colonization^12^. However, motility regulation overlaps with a complex signaling network that controls functions involved in biofilm formation such as quorum sensing or exopolysaccharide production^13,14^. It has therefore been difficult to determine both a clear causal relationship and the mechanisms that could support a hypothesis of motility as an advantageous feature for biofilm-forming bacteria.

In *E. coli,* flagellar activity has been proposed to facilitate the initial contact of the cell with the surface, potentially helping to overcome repulsive forces on the surface^10^. Nevertheless, when other surface appendages such as Curli or conjugative pili are constitutively expressed, flagella become dispensable for the initial adhesion and biofilm development^15,16^, suggesting motility *per se* might not be the adhesion promotion factor. On the other hand, flagella mechanosensory function as a surface-sensing tool has been proposed to govern the planktonic-sessile transition underlying biofilm formation^17,18^. However, in this case the surface detection by the flagella culminates in motility downregulation. Consistently, high concentrations of the second messenger cyclic diguanylate (c-di-GMP) have been shown to correlate with motility downregulation and the development of a thick biofilm^19,20^. These results seem paradoxical with regard to the positive effect of motility on biofilm development, highlighting motility and biofilm development as mutually exclusive events. Nevertheless, bacteria may also swim within a mature biofilm^21^.

Motility has also been suggested to influence biofilm maturation and architecture^11,22^, although reports about the impact of cell motility on later stages of biofilm development are scarce. In *Vibrio cholerae,* motility has been proposed to favor the invasion of resident biofilms^23^. Ultimately, the motility effect on biofilm formation significantly changes depending on the environment, including surface properties and hydrodynamics^24^. Therefore, despite its obvious competitive fitness advantage in planktonic life (particularly regarding nutrient pursuit), the question of whether cell motility is a superior trait in surface-colonizing competition remains open.

To address this issue, we examined biofilm formation by motile and nonmotile *E. coli* cells in the controlled geometry of a millifluidic device from a kinetic perspective, covering both the short and long time scales of the adherent community development. This allowed us to search for mechanistic information that could distinguish the colonizing ability of swimming cells from their nonmotile counterparts. In this study, we take motility to mean flagellar motility apart from surface-associated motions such as swarming or twitching^25^. We thus examined the simple situation of motile vs. nonmotile *E. coli* strains colonizing a bare glass surface, followed by a more complex and more naturally relevant situation involving the same strains in the presence of other species in a co-colonization test. The latter included a 4- species assemblage that we previously showed to form a deterministic community in about 40 hours of growth under continuous nutrient flow in a millifluidic channel^26^. The use of this continuous flow growth mode makes it possible to control the physicochemical properties of the environment throughout the development of the biofilm, and ensures that the biofilms can be thoroughly compared.

Our results reveal a clear-cut effect of motility on surface colonization, which consists in introducing a lag time of several hours to the biofilm development. Meanwhile, the motile and nonmotile cells display a similar biofilm growth rate. We interpret this effect in terms of a spatial exploration discrepancy dominated by characteristic times of settling and diffusion, using a first passage mathematical model. Finally, we reveal how the presence of co-colonizers affects this behavior and discuss the strong dependence of flagellar motility effects on specific environmental conditions.

## Materials and Methods

### Bacterial strains and culture conditions

*E. coli* strains were derived from *E. coli* K-12 classified as MG1655. In addition to the wild type motile strain, we used the nonmotile variant which lacks the insertion sequence IS*1* upstream of the *flhD* promoter. This consequently blocks cell swimming, as shown by flagellar motility tests (Fig. S1). Both strains express the conjugative F-pilus carried by a plasmid, which ensures robust biofilm formation^27^, and the FAST-mCherry fusion protein to provide biofilm-relevant fluorescence labeling^28^. For surface dynamics monitoring experiments, we used variants that constitutively express GFP. The co-colonizers belong to a previously described four-species community^29^ consisting of *Bacillus thuringiensis* (*Bt*), a 407 Cry^-^ strain, *Pseudomonas fluorescens (Pf)* (WCS365), *Kocuria varians (Kv)* (CCL56), and *Rhodocyclus* sp. *(Rh)* (CCL5). For 4S biofilm kinetic monitoring, we used fluorescent *Bt* and *Pf* variants carrying the FAST gene on the chromosome^26^ and plasmidic mCherry (pMP7605)^30^, respectively. The strains were routinely cultivated at 30°C on M1 medium (see Supplementary Information).

### Millifluidic device

Millifluidic channels were microfabricated to be 30 mm x 1 mm x 1 mm (length x width x height). A polydimethylsiloxane (PDMS) mixture (RTV615A+B; Momentive Performance Materials) was poured at ambient temperature in a polyvinyl chloride home-micromachined mold and left to cure at least 3 hours in an oven set at 65°C. Next, the recovered templates were drilled for further plugging of adapted connectors and tubing. PDMS templates and glass coverslips were then cleaned using an oxygen plasma cleaner (Harrick) and immediately bound together to seal the channels. For connections, we used stainless steel connectors (0.013” ID and 0.025” OD) and microbore Tygon tubing (0.020” ID and 0.06” OD) supplied by Phymep (France). The thin metallic connectors provide a bottleneck in the flow circuit, which prevents upstream colonization. The medium was pushed into the channels at a rate of 1 ml/h with syringe pumps for the 36-40 hours of the experiment. The whole experiment was thermostatically maintained at 30°C.

### Biofilm formation

*Initiation:* 1.2×10^5^ cells were obtained from exponentially growing cultures. Cell injections were performed directly into the PDMS channels using a syringe equipped with a 22G needle before connecting the tubing. Next, the cells were allowed to settle for 1h30 before starting the medium flow. All times at t=0 referred to the flow triggering time. For biofilm growth, we used MB medium, which is adapted from M1 medium (details provided in Supplementary Information). Overnight cultures in M1 — seeded with a single colony from M1-agar plates — were grown at 30°C under agitation. Exponential phases were obtained from dilutions in M1 of these overnight cultures incubated at 30°C under agitation. The same protocol was applied in the presence of co-colonizers, except that 1.2×10^5^ cells from exponentially growing cultures of each co-colonizer were injected at the same time as *E. coli* into the channel.

### Microscope imaging

*Microscopy:* We used an inverted NIKON TE300 microscope equipped with motorized x, y, and z displacements and shutters. Images were collected using a 20’ S plan Fluor objective (NA 0.45 WD 8.2-6.9 mm). Bright-field images were collected in direct illumination (no phase). Fluorescence acquisitions were performed using either the green channel filters for GFP and FAST:HBR-2,5-DM (Ex. 482/35, DM 506 Em. FF01-536/40) or the red filter for m- Cherry (Ex 562/40nm DM 593 Em. 641/75). Excitation was performed using an LED box (CoolLed pE-4000). For dynamics measurements, confocal images were collected using a spinning disk Crest X light V2 module (Gataca, France distribution) with an axial resolution of 5.8 μm.

### Image acquisition

A Hamamatsu ORCA-R2 EMCCD camera was used for time-lapse acquisitions of 1344×1024 pixel images with 12-bit grey level depth (4096 grey levels), and to capture an *xy* field of view of 330 μm’ 430 μm. Bright-field and fluorescence images were typically collected for 36 hours at a frequency of 6 frames per hour. Excitation LEDs were set at a 50% power level, and exposure times were 50 ms or 500 ms for the green emissions (for GFP and FAST, respectively) and 800 ms for the red emissions.

### Image analysis

*Image intensities:* Time-lapse images were analyzed to derive the kinetics of *E. coli* biomass accumulation in the channel based on FAST fluorescence intensity, as previously detailed^28^. Image intensity per pixel, averaged for the whole image or on defined regions of interest (ROIs), was collected using the NIKON proprietary software NIS. Subsequently, the data sheets edited by NIS were exported to MATLAB for further analysis of biofilm development kinetics. Background was subtracted using the contribution to the fluorescence intensity of a channel containing medium without any bacteria. All curves were averaged over at least three independent positions and two independent replicates.

### Dynamics monitoring

The individual dynamics of *E. coli* cells were tracked in the biofilm surface layer using confocal time-lapse acquisitions performed with a 60x, 1.4NA objective. Series of 50 images were recorded at an acquisition frequency of 90 frames per hour every 2 hours, for 24 h. The image stacks were binarized using the ImageJ IsoData threshold calculation tool, which iteratively takes into account average background and average object intensities. Next, objects comprising between 40-2,000 pixels in area (1px=0.1075 μm) were tracked in 2D over each temporal series using the ImageJ MTrack2 algorithm. These size limits ensured the tracking of a single cell to clusters of a few (3-4) cells. The minimal trajectory held 2 frames and the maximal accepted displacement between two frames was 60 pixels. Thereafter, the results file containing the sorted coordinates of all the trajectories was exported to MATLAB in order to calculate the trajectory persistence *P.* For each time series, *P* was obtained by taking the average over all the trajectories *i* of the individual persistence *P_i_*, defined as the ratio of the travelled distance *d_i_* to the trajectory length *L_i_*.

## Results

### Motility strongly delays bare surface colonization by *E. coli*

We assessed how fast the bacteria form a biofilm under mild flow depending on their motility by comparing the surface colonization kinetics of the nonmotile cells (Mot^-^) with that of their swimming counterparts (Mot^+^). Time-lapse images were taken from the very beginning of the process, consisting of a few adhering cells, up to the stage of a dense cell material after 48 hours of growth. The results show very distinct kinetic profiles for motile and nonmotile cells (Fig. 1A). Using a logarithm scale to display the fluorescence intensity as a function of time revealed that this difference mainly consists in a long lag time of about 20 hours, whereas almost no lag time was evident for nonmotile cells (Fig. 1B). In contrast, the colonization rate measured after the lag appears very similar for motile and nonmotile bacteria, indicating that the biofilm initiation phase was essentially altered when cells were motile as opposed to nonmotile. Furthermore, we measured the cell division rates under planktonic growth in MB medium and observed no differences (Fig. S2). To determine the details of this initial phase, we recorded stacks of images at a higher frequency (90 images per hour instead of 6) every two hours, using a higher magnification objective (63x) and confocal acquisitions in order to make observations at the single cell level and evaluate the surface dynamics. Applying a basic 2D tracking routine to this image series, we collected cell trajectories for each time series and derived a persistence index, *P*. The mean of all individual trajectory *P_i_* values was defined as the ratio of the trajectory length *L_i_* to the traveled distance *d_i_* as shown in Fig. 2A. This index provides a quantitative evaluation of the cell dynamics on the surface that varies from 0 for a fully steady cell to 1, which corresponds to the highest displacement in a straight line authorized in the analysis, i.e. the strongest dynamics. The analysis concentrates on the initial phase of the colonization where cell population is limited enough to enable single cell delineation, corresponding to the first 6 hours of biofilm formation for nonmotile cells and extending up to 24 hours for motile cells. The results in Fig. 2 (panels B and C) indicate that motile and nonmotile cells exhibit trajectories with similar persistence on the surface. In some cases, motile cells even counterintuitively displayed a smaller average persistence than nonmotile cells, particularly in the first hours of the biofilm formation. However, it should be noted that only a few events contributed to the determination of *P* for motile cells in this time period, which increased the weight of potentially fixed individuals in the calculation. Comparing equivalent colonization degrees, very close persistence values were obtained for time t=4h (Fig. 2C). The small number of cells on the surface also explains the high observed standard deviation of the persistence of motile cells.

**Figure 1:**
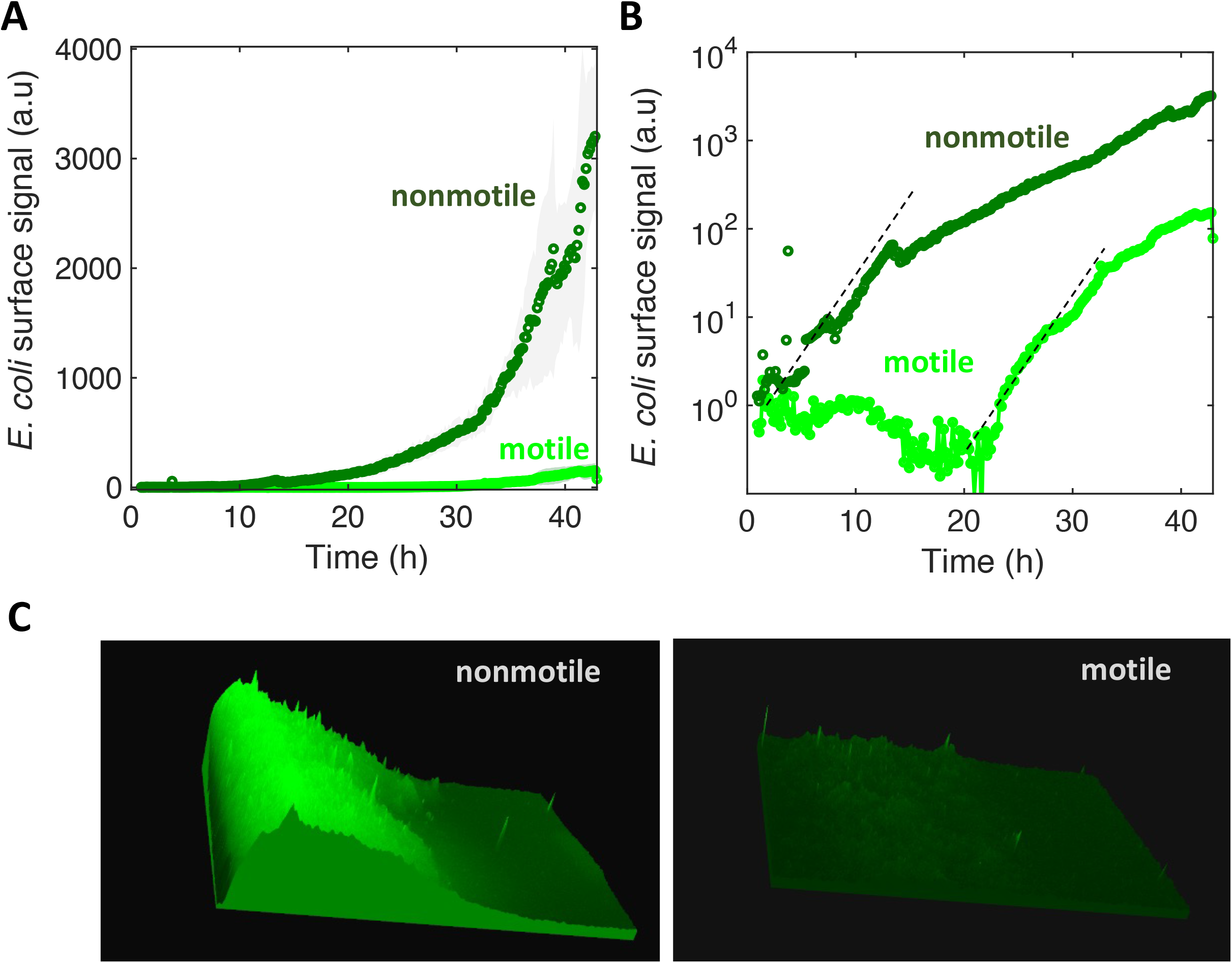
Motility delays biofilm formation. (A) *E. coli* FAST fluorescence intensity as a function of flow time for a biofilm grown by motile (bright green) and nonmotile (dark green) cells in a millifluidic channel; bold curves are from the average intensity of 3 distinct positions in at least 2 different channels with the shaded area representing the standard deviation (B) Same data as in A except the logarithm of the average fluorescent intensity is plotted. The biofilm was grown at a rate of 1 ml/h medium flow, at 30°C. Dashed lines highlight the exponential part of the growth and display the same slope for both motile and nonmotile cells.(C) Fluorescence intensity surface plots from biofilm images recorded at time t=30h. Biofilm from nonmotile cells on the left panel and from motile on the right panel.

**Figure 2:**
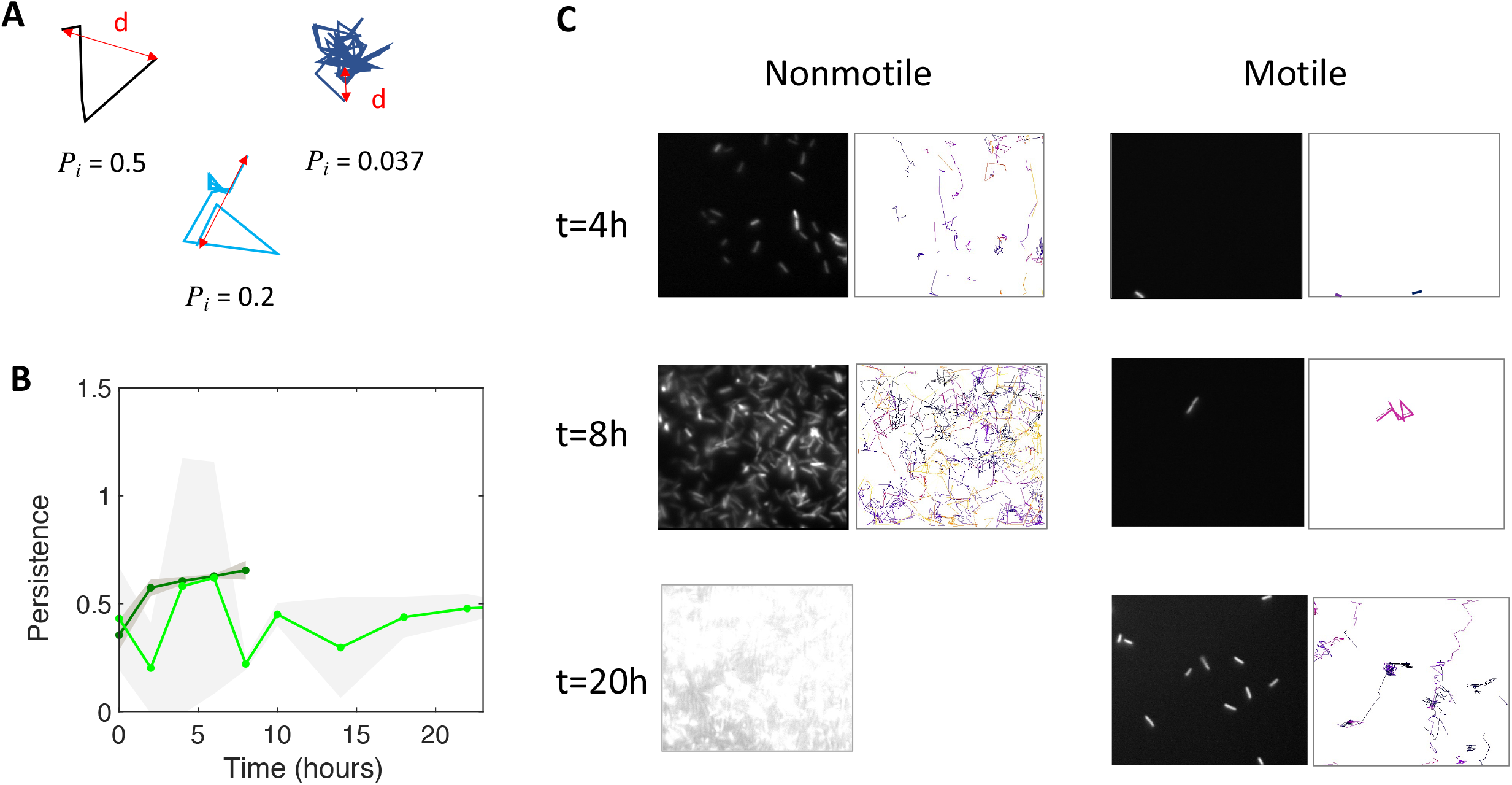
Motile and nonmotile cells display similar surface persistence in the biofilm initiation phase. (A) Typical trajectories with their respective persistence index values. (B) Persistence values over time for motile (light green) and nonmotile (dark green) cells on the surface. The curves represent the average, with the shaded area representing the standard deviation. (C) Snapshots of the biofilm development together with detected trajectories for nonmotile (left columns) and motile cells (right columns). At t=20h, the surface of nonmotile cells was overcrowded, which excludes the ability to track single cells.

The fraction of cells reaching the surface, and not their dynamics on the surface, is therefore more likely to explain the lingering of motile cells during biofilm initiation as compared to Mot^-^ cells. To account for these results, we examined whether cell bulk behavior could be an important driving force of colonization. To formalize this idea, a simple theoretical model was created to describe motile versus nonmotile cell behavior.

### Motility increases the characteristic time for cell-surface interaction

In our model, we considered a bacterial suspension in a box (defined by volume *V* and height *H*) and we wrote equations to describe the variation with time for the number of cells reaching the bottom surface upon settling (nonmotile cells) as well as upon run-and-tumble diffusion (motile cells). As detailed in the Supplementary Information, settling of nonmotile cells occurred according to the unidirectional speed Vs (Stokes’ law) of a particle submitted to gravity in a viscous medium. This therefore provides a number of cells 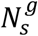 on the surface as a function of time (for *t* < *H/V_s_*):

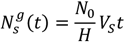

(For *t* > *H/V_s_*, all cells have reached the bottom surface and 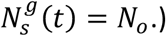

For diffusing cells (with diffusion coefficient *D*), we calculated the kinetics of the first passage to an absorbing boundary using the universal equation of Meyers and colleagues (details provided in the Supplementary Information) and determined 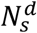, the number of cells reaching the surface as a function of time:

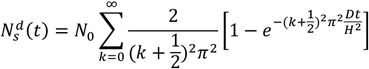

The results displayed in Fig. 3 show that the time for 99% of the population to reach the surface by settling is 1h30, whereas the same proportion of swimming bacteria with a diffusion coefficient of 10 μm^2^/s require 52 hours to make their first passage on the surface^31^. These findings are fully consistent with the experimental observations. Starting with the same number of motile and nonmotile cells and by making a reasonable hypothesis about biofilm development kinetics following a logistic law that closely fits the data (Supplementary Information, Fig. S3), we find that the motile-nonmotile cell asymmetry in the cell number on the surface explains the observed significant delay in biofilm growth. Indeed, as flow begins, the non-attached cells are removed from the channel and no longer participate in biofilm formation. This means that determining motility function consists in modulating the number of cells that reach the surface per unit of time. Our model also predicts the impact of both the device geometry and the diffusion coefficient on the surface rate (Supplementary Information, Fig. S4). We found that swimming increasingly delays surface access when the channel height is greater than 20 μm. In addition, simulations show that surface access delay increases as swimming rates (i.e. the diffusion coefficient) decrease (Supplementary Information, Fig. S5).

**Figure 3:**
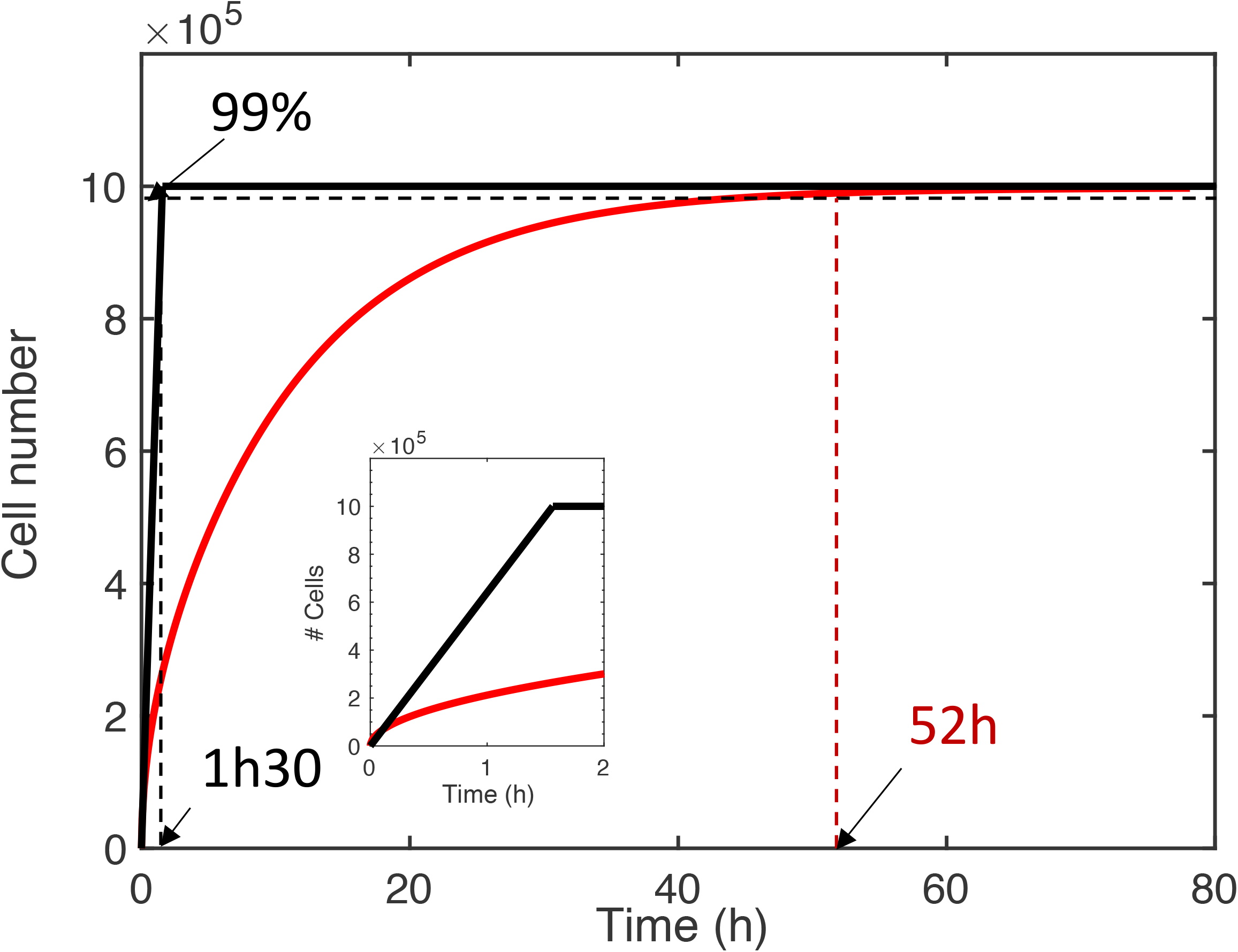
Calculated kinetics of settling (motile cells) and diffusion (nonmotile cells) towards the surface. The number of cells reaching the surface as a function of time in a 1-mm high channel was calculated according to settling (in black) and diffusion (in red) equations 1 and 2, using the diffusion coefficient D=10 μm^2^/s; cell size a=1 μm; g=10 m/s^2^; and cell-medium mass density contrast *Δρ* = 80 *kg/m*^3^. The black dashed line indicates the end of the incubation time and the start of the flow at which point 99% of the cells have touched down by settling; the red line indicates the time for 99% of the motile cells to touch down by diffusion. The inset shows a zoom in on the two first hours.

Altogether, these results suggest that surface access, and not surface attachment once the cell has reached the surface, is the limiting step of the process, under the conditions used in this study.

### The presence of co-colonizers affects the surface colonization process for both motile and nonmotile cells

In order to evaluate if cell motility brings about a similar lag time in a more complex environment, we monitored how motile and nonmotile *E. coli* cells colonize a bare surface in the presence of co-colonizers. For this, we examined surface colonization by *E. coli* in the presence of four other strains that we recently showed are able to deterministically form a stable biofilm under flow in about 40 hours^26^.

The four species were introduced in the channel at the same time as the motile or nonmotile *E. coli* cells, following the same procedure used for single species biofilm formation and kinetics monitoring. The results in Fig. 4 (5S) show that motile bacteria still display a slight deficit in colonization efficiency, although the difference between motile and nonmotile cells was significantly reduced in the presence of the co-colonizers. Notably, the presence of the co-colonizers altered not only the amplitude but also the kinetic phases of development, as highlighted by the logarithmic display of the curves (Fig. 4B). In the presence of the co-colonizers, we observed the succession of two phases displaying the same timing for motile and nonmotile bacteria. The first phase consisted of an initial limited surface growth that leveled about 6 hours after starting the nutrient flow, with a plateau that extended approximately up to 12 to 15 hours into colonization. Subsequently, a second growth phase started which increased the *E. coli* biomass in the composite biofilm by 2 logs. The curves indicate that the motile cell biofilm entered the second growth phase with a small 2- to 3-hour delay.

**Figure 4:**
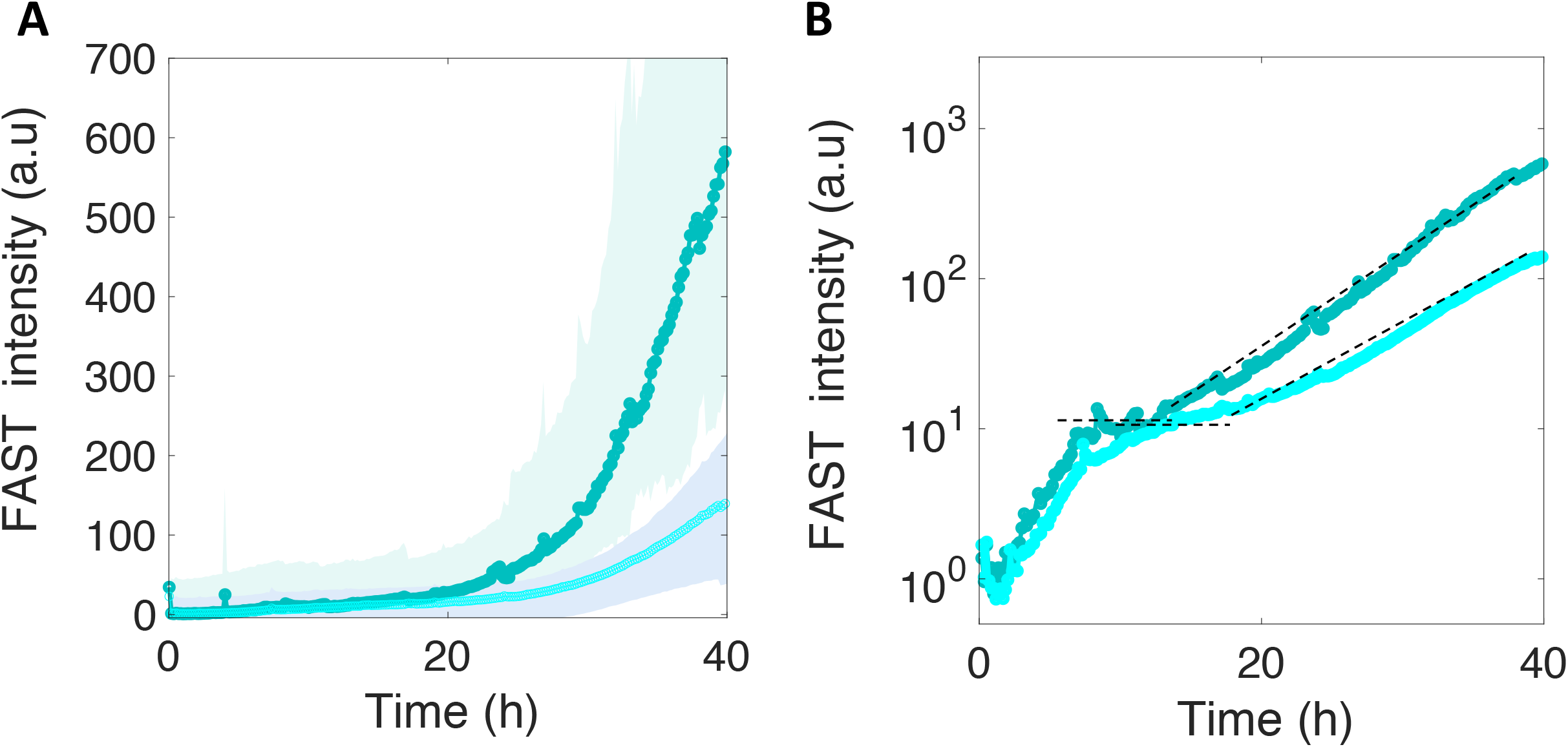
Motile and nonmotile *E. coli* surface colonization in the presence of co-colonizers. Motile (in light cyan) and nonmotile (in dark cyan) *E. coli* surface fluorescence (A) in the biofilm formed in the presence of the four co-colonizers *Bt, Pf, Kv* and *Rh.* The logarithmic display (B) highlights two distinct phases in *E. coli* development in the composite multispecies biofilm. The dashed lines indicate the level of the first phase plateau and the second phase growth rate.

We next asked how these kinetics could be related to the development of the biofilm colonizers. To answer this question, we monitored the development of the four-species biofilm in parallel channels, after having introduced fluorescent variants of *Bacillus thuringiensis* (*Bt-*FAST) and *Pseudomonas fluorescens* (*Pf*-mCherry), the two main contributors in the four-species community. Fig. 5A-B specifically shows the kinetics of these two fluorescent species in the four-species biofilm.

**Figure 5:**
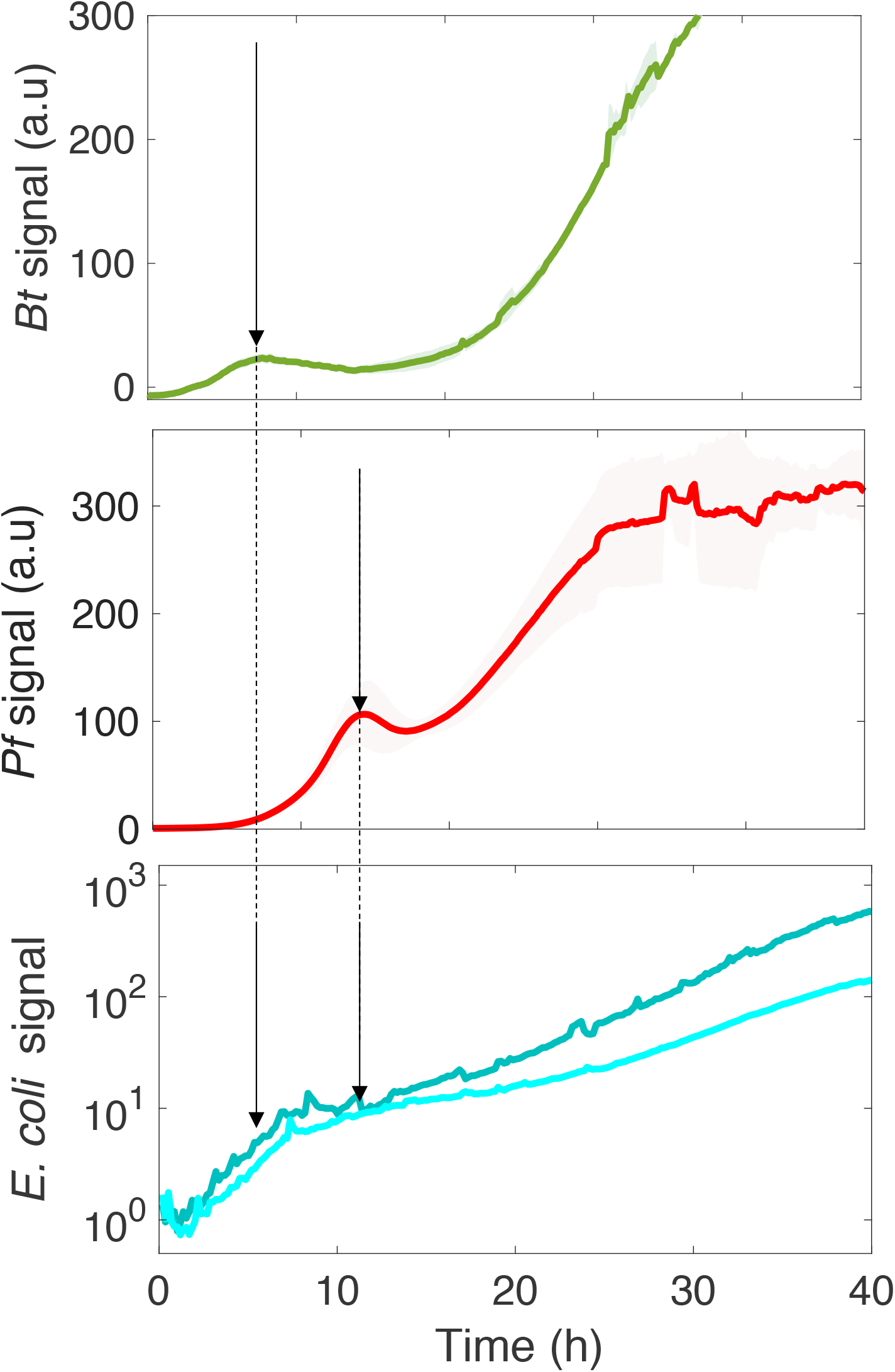
The kinetics of motile and nonmotile *E. coli* surface colonization follows the community development timing of the four species. *Bt* (A) and *Pf* (B) surface development kinetics in the four-species community. In parallel, the logarithmic display of the *E. coli* signal on the surface is shown in the presence of the co-colonizers (C). The arrows point to the *Bt* (A) and *Pf* (B) initial climaxes, and the *E. coli* development phases (C).

The saturation of *E. coli* development in its initial phase coincided with the first climax of *Bt* in the four-species community, whereas the second growth phase of *E. coli* started as *Pf* development reached its first climax. These results indicate that the deterministic biofilm formation program of the co-colonizers dominated the *E. coli* surface colonization process, inducing a kinetic remodeling and almost completely abolishing the difference between motile and nonmotile cells in their colonization efficiency. This indicates that the limiting step of the surface colonization by *E. coli* is shifted by the presence of the co-colonizers, significantly altering the impact of motility in the process.

Finally, we also tested the colonization of the pre-established four-species biofilm by the motile and nonmotile strains of *E. coli* at 8, 20 and 36 hours after inoculation and incubation of the four species. This demonstrated that the corresponding signals barely emerged from the background for both motile and nonmotile cells (Supplementary Information), indicating that neither of the two strains were able to colonize the surface in the presence of the pre-established strains.

## Discussion

Bacterial flagellar motility has been investigated for decades now, and significant advances in the understanding of the involved mechanisms have been made. However, many open questions remain about the impact of this function on the ability of bacteria to colonize surfaces. Here, we report a kinetic analysis of *E. coli* surface colonization in a millifluidic channel, with a focus on active vs. non-active flagellar motility. We found that the impact of motility on the biofilm-forming capacity is highly dependent on both the topology and the populations of the environment. Our results show that motility introduces a significant delay to the colonization of bare surfaces by *E. coli* alone. Based on mathematical modeling of the settling and diffusion of the bacteria in the channel, we were able to simulate the respective kinetics of surface landing and demonstrate that bacterial motility induces a significant increase in the mean characteristic time needed for the whole cell population to reach the surface in a millimetric geometry. This is in contrast to the intuitive concept encountered in the literature that the absence of motility reduces the chances of bacterial cells coming into contact with the surface^10,24^. In our experimental configuration, the ‘race’ towards the surface is dominated by the cell axial velocity which is higher for settling bacteria. In contrast, instantaneous velocity is higher for motile bacteria but does not favor access to the surface.

The analytic expressions that we derived for settling and diffusion highlights the importance of the geometry of the environment, which has generally been overlooked in colonization assays. This shows that competition between motile and nonmotile cells depends on the height of the environmental compartment. Indeed, we found that motile cells are more successful in shorter-walled channels than taller ones. Our simple model predicts a crossover at a height of 100 μm, at which point diffusion takes over settling and the competitive advantage of being the first on the surface shifts to motile cells. The channel geometry that we used is relevant for many natural situations in which bacteria dwell in channels and pores in the millimetric range under continuous or intermittent flow.

We observed that local dynamics and biomass growth rates were very similar for both motile and nonmotile cells, suggesting that flagellar motility does not alter anchorage or surface persistence once the cells have reached the surface, an issue that has remained controversial to date. As detailed in previous investigations of this question^10^, it is difficult to decipher the primary factors involved in the initial attachment. A particular challenge is to disentangle the contribution of motility *per se* from the contribution of flagella as a surface appendage and the contingent involvement of other structures such as type I pili. Under the conditions used in this study for bare surface colonization, it is likely that the attachment step is dominated by the overexpression of the F-pilus, which is crucial in promoting the initial adhesion^15,27,32^. In this case, flagellar motility simply reduces surface abundance, which is established during the inoculation period, and does not affect biofilm development. Mathematical model predictions can also demonstrate the impact of different diffusion coefficients on surface abundance over time. This is an interesting finding to take into account for multispecies colonization processes where surface access kinetics are crucial in the competitive dynamics that shape the attached community^33^.

In the presence of co-colonizers (here, the members of a 4-species community able to build a stable biofilm), we observed that motile and nonmotile *E. coli* cells exhibit very similar colonization profiles that differ from the profiles displayed in the single-species colonization experiments. Specifically, there is a lower surface-bound *E. coli* global biomass, consistent with the intrinsic competition for the surface expected from the presence of other adhesive species^34^. Two striking features stand out here: the emergence of *E. coli* kinetic colonization phases that match the four-species biofilm climaxes; and the reduced lag observed between motile and nonmotile cells in the characteristic time of colonization, primarily due to the receding of the nonmotile cells in comparison to the *E. coli* colonizing ability in the absence of co-colonizers.

In a previous study, the four-species biofilm climaxes were interpreted as species responses to the oxygen depletion induced by biofilm development, suggesting that the reduction in oxygen might also contribute to the diminished colonizing efficiency of *E. coli*^26^. Moreover, knowing that the lack of oxygen strongly affects *E. coli* motility^35^ by inducing a motile to nonmotile transition in the bacterial population, we can hypothesize that the environmental oxygen scarcity caused by the co-colonizers accounts for the convergence of the motile and nonmotile cell colonization kinetics in this multispecies context. Nevertheless, the co-colonizers could also induce a shift in the limiting step of the colonizing process by increasing the characteristic time of attachment on the surface, which would result in abolishing the difference between motile and nonmotile cells that ultimately dwell on the surface. These results stress the importance of studying the processes from a kinetic perspective in order to acquire mechanistic information.

Our report establishes that the impact of flagellar motility on surface colonization is not an intrinsic trait associated with this function; instead, it closely depends on an environment defined by both topology and population composition. Importantly, we demonstrate that cell swimming essentially regulates the surface access time. However, a shift in the environmental conditions (such as the presence of co-colonizers) can drastically alter the outcome of this distinctive property and abolish the asymmetry between motile and nonmotile cells. We thus propose here a model with the potential to resolve the long-standing controversy over the role of cell motility in surface colonization and biofilm formation.

## Supporting information

https://docs.google.com/document/d/1xHXgqMGN5M0pxBk9dVIUo_obkDSGFr8M/edit

## Acknowledgments

The authors would like to thank Rapahaël Voituriez and Philippe Thomen for their fruitful discussions, and Carounagarane Doré for technical assistance in developing the devices.

This work was supported by a MESRI fellowship to WBY.

## References

1 Flemming, H. C. et al. Biofilms: an emergent form of bacterial life. Nat Rev Microbiol 14, 563–575, doi:10.1038/nrmicro.2016.94 (2016).

2 Hall-Stoodley, L., Costerton, J. W. & Stoodley, P. Bacterial biofilms: from the natural environment to infectious diseases. Nat Rev Microbiol 2, 95–108, doi:10.1038/nrmicro821 (2004).

3 Karygianni, L., Ren, Z., Koo, H. & Thurnheer, T. Biofilm Matrixome: Extracellular Components in Structured Microbial Communities. Trends Microbiol 28, 668–681, doi:10.1016/j.tim.2020.03.016 (2020).

4 Berg, H. C. & Anderson, R. A. Bacteria swim by rotating their flagellar filaments. Nature 245, 380–382, doi:10.1038/245380a0 (1973).

5 Silverman, M. & Simon, M. Flagellar rotation and the mechanism of bacterial motility. Nature 249, 73–74, doi:10.1038/249073a0 (1974).

6 Nakamura, S. & Minamino, T. Flagella-Driven Motility of Bacteria. Biomolecules 9, doi:10.3390/biom9070279 (2019).

7 Colin, R., Ni, B., Laganenka, L. & Sourjik, V. Multiple functions of flagellar motility and chemotaxis in bacterial physiology. FEMS Microbiol Rev, doi:10.1093/femsre/fuab038 (2021).

8 O’Toole, G. A. & Kolter, R. Flagellar and twitching motility are necessary for Pseudomonas aeruginosa biofilm development. Mol Microbiol 30, 295–304, doi:10.1046/j.1365-2958.1998.01062.x (1998).

9 Lemon, K. P., Higgins, D. E. & Kolter, R. Flagellar motility is critical for Listeria monocytogenes biofilm formation. J Bacteriol 189, 4418–4424, doi:10.1128/JB.01967-06 (2007).

10 Pratt, L. A. & Kolter, R. Genetic analysis of Escherichia coli biofilm formation: roles of flagella, motility, chemotaxis and type I pili. Mol Microbiol 30, 285–293, doi:10.1046/j.1365-2958.1998.01061.x (1998).

11 Wood, T. K., Gonzalez Barrios, A. F., Herzberg, M. & Lee, J. Motility influences biofilm architecture in Escherichia coli. Appl Microbiol Biotechnol 72, 361–367, doi:10.1007/s00253-005-0263-8 (2006).

12 Josenhans, C. & Suerbaum, S. The role of motility as a virulence factor in bacteria. Int J Med Microbiol 291, 605–614, doi:10.1078/1438-4221-00173 (2002).

13 Merritt, P. M., Danhorn, T. & Fuqua, C. Motility and chemotaxis in Agrobacterium tumefaciens surface attachment and biofilm formation. J Bacteriol 189, 8005–8014, doi:10.1128/JB.00566-07 (2007).

14 Shrout, J. D., Tolker-Nielsen, T., Givskov, M. & Parsek, M. R. The contribution of cell–cell signaling and motility to bacterial biofilm formation. MRS Bull 36, 367–373, doi:10.1557/mrs.2011.67 (2011).

15 Reisner, A., Haagensen, J. A., Schembri, M. A., Zechner, E. L. & Molin, S. Development and maturation of Escherichia coli K-12 biofilms. Mol Microbiol 48, 933–946, doi:10.1046/j.1365-2958.2003.03490.x (2003).

16 Prigent-Combaret, C. et al. Developmental pathway for biofilm formation in curli-producing *Escherichia coli* strains: role of flagella, curli and colanic acid. Environ Microbiol 2, 450–464 (2000).

17 Wong, G. C. L. et al. Roadmap on emerging concepts in the physical biology of bacterial biofilms: from surface sensing to community formation. Phys Biol 18, doi:10.1088/1478-3975/abdc0e (2021).

18 Laganenka, L., Lopez, M. E., Colin, R. & Sourjik, V. Flagellum-Mediated Mechanosensing and RflP Control Motility State of Pathogenic Escherichia coli. mBio 11, doi:10.1128/mBio.02269-19 (2020).

19 Valentini, M. & Filloux, A. Biofilms and Cyclic di-GMP (c-di-GMP) Signaling: Lessons from Pseudomonas aeruginosa and Other Bacteria. J Biol Chem 291, 12547–12555, doi:10.1074/jbc.R115.711507 (2016).

20 Jenal, U., Reinders, A. & Lori, C. Cyclic di-GMP: second messenger extraordinaire. Nat Rev Microbiol 15, 271–284, doi:10.1038/nrmicro.2016.190 (2017).

21 Houry, A. et al. Bacterial swimmers that infiltrate and take over the biofilm matrix. Proc Natl Acad Sci U S A 109, 13088–13093, doi:10.1073/pnas.1200791109 (2012).

22 Barken, K. B. et al. Roles of type IV pili, flagellum-mediated motility and extracellular DNA in the formation of mature multicellular structures in Pseudomonas aeruginosa biofilms. Environ Microbiol 10, 2331–2343, doi:10.1111/j.1462-2920.2008.01658.x (2008).

23 Nadell, C. D., Drescher, K., Wingreen, N. S. & Bassler, B. L. Extracellular matrix structure governs invasion resistance in bacterial biofilms. ISME J 9, 1700–1709, doi:10.1038/ismej.2014.246 (2015).

24 Zheng, S. et al. Implication of Surface Properties, Bacterial Motility, and Hydrodynamic Conditions on Bacterial Surface Sensing and Their Initial Adhesion. Front Bioeng Biotechnol 9, 643722, doi:10.3389/fbioe.2021.643722 (2021).

25 Wadhwa, N. & Berg, H. C. Bacterial motility: machinery and mechanisms. Nat Rev Microbiol, doi:10.1038/s41579-021-00626-4 (2021).

26 Monmeyran, A. et al. Four species of bacteria deterministically assemble to form a stable biofilm in a millifluidic channel. NPJ Biofilms Microbiomes 7, 64, doi:10.1038/s41522-021-00233-4 (2021).

27 Ghigo, J. M. Natural conjugative plasmids induce bacterial biofilm development. Nature 412, 442–445 (2001).

28 Monmeyran, A. et al. The inducible chemical-genetic fluorescent marker FAST outperforms classical fluorescent proteins in the quantitative reporting of bacterial biofilm dynamics. Sci Rep 8, 10336, doi:10.1038/s41598-018-28643-z (2018).

29 Sheppard, A. E., Poehlein, A., Rosenstiel, P., Liesegang, H. & Schulenburg, H. Complete Genome Sequence of Bacillus thuringiensis Strain 407 Cry. Genome Announc 1, doi:10.1128/genomeA.00158-12 (2013).

30 Lagendijk, E. L., Validov, S., Lamers, G. E., de Weert, S. & Bloemberg, G. V. Genetic tools for tagging Gram-negative bacteria with mCherry for visualization in vitro and in natural habitats, biofilm and pathogenicity studies. FEMS Microbiol Lett 305, 81–90, doi:FML1916 [pii] 10.1111/j.1574-6968.2010.01916.x (2010).

31 Patteson, A. E., Gopinath, A., Goulian, M. & Arratia, P. E. Running and tumbling with E. coli in polymeric solutions. Sci Rep 5, 15761, doi:10.1038/srep15761 (2015).

32 Beloin, C., Houry, A., Froment, M., Ghigo, J. M. & Henry, N. A short-time scale colloidal system reveals early bacterial adhesion dynamics. PLoS biology 6, e167 (2008).

33 Eigentler, L. et al. Founder cell configuration drives competitive outcome within colony biofilms. ISME J, doi:10.1038/s41396-022-01198-8 (2022).

34 Lloyd, D. P. & Allen, R. J. Competition for space during bacterial colonization of a surface. J R Soc Interface 12, 0608, doi:10.1098/rsif.2015.0608 (2015).

35 Douarche, C., Buguin, A., Salman, H. & Libchaber, A. E. Coli and oxygen: a motility transition. Phys Rev Lett 102, 198101 (2009).

