## Supplementary material for "Flagellar Motility During E. Coli Biofilm Formation Provides a Competitive Disadvantage Which Recedes in the Presence of Co-Colonizers": https://docs.google.com/document/d/1xHXgqMGN5M0pxBk9dVIUo_obkDSGFr8M/edit

### Supplementary information

#### I. Materials and methods

- **Media composition**

| *Composition* | **M1** (g/L) | **MB** (g/L) |
|---|---|---|
| Yeast Nitrogen Base* | 1.7 | 1.7 |
| Ammonium sulfate | 5 | 5 |
| Glucose | 10 | 0.4 |
| Casamino-acids | 5 | 1 |

*from DIFCO BD

- **Motility assay**

Motile and nonmotile strains swimming motility was tested according to an adaptation of the protocol of Barker and collaborators [1]. Aliquots of the same concentration ($10^6$cells/ml) of motile and nonmotile *E. coli* cells from overnight cultures were inoculated into a soft agar (LB semi-solidified with 1.25% wt/vol agar enabling flagellar motility [1]. The plates were incubated at 30°C during 24h and imaged. The results are shown in Fig. S1. The diffuse spreading of the motile cells can be easily recognized by eye in comparison with the small nonmotile cells colony where no outgrowth occurred.

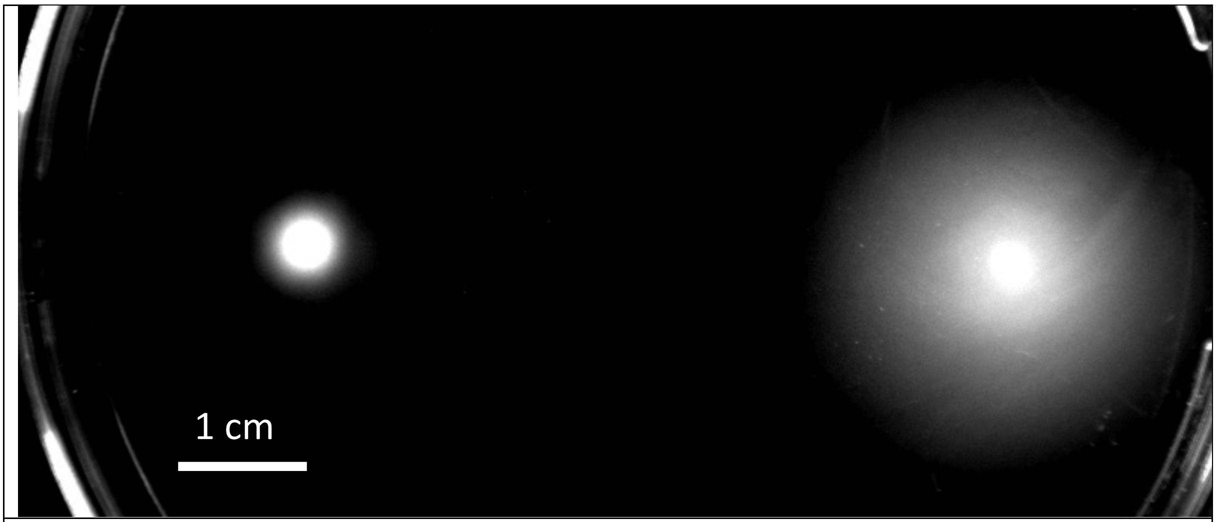


**Figure S1: Motility assay**. Image of nonmotile (on the left) and motile (on the right) cells inoculated in 1.25% wt/vol agar after 24 hours growth in MB medium.

- **Motile and nonmotile cells exhibit similar growth rates**

In order to detect potential difference in the division rates of motile and nonmotile cells, we measured both strains optical densities over time in MB medium at 30°C using a microplate reader (Tecan TECAN Infinite M200 pro equipped with UV Xenon flashlamp light source). $10^6$ cells ($10^6$cells/ml exponentially growing) were seeded in each well in triplicate and left to grow over night, taking one measurement every 10 min. The curves displayed in Fig. S2 show no significant difference between motile and nonmotile growth.

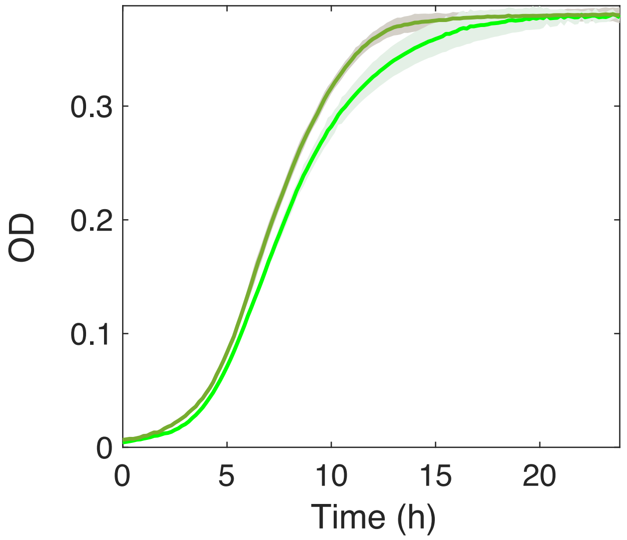


**Figure S2. Motile and nonmotile cells have similar division rates under planktonic growth.** Nonmotile cells (dark green) and motile cells (light green). $10^6$ cells/ml were seeded in MB medium. Measurements performed in the triplicates in 48-wells plates. Curves represent the triplicate average shaded with the standard deviation of the data set.

#### II. Mathematical model description

We aim to predict the kinetics of adhesion to the bottom side of a chamber for a suspension of bacteria, in two cases: motile bacteria and non-motile bacteria. The chamber volume is V (height H and bottom surface S, V=SH). We assume:

- Adhesion to the bottom surface is immediate at first contact.
- No hydrodynamic effect when a cell approaches the bottom surface (no trapping, no bouncing).
- Initially, the spatial distribution of bacteria across the chamber is uniform.

For the sake of simplicity, the problem is restricted to 1D.

**a. Non-motile cells:**

Stokes' law gives the terminal settling speed of a round particle in a viscous fluid:

$$V_s = \frac{2\,a^2 g\Delta\rho}{9\eta}$$

with $a$ the particle radius, $g$ is the gravitational acceleration, $\Delta\rho$ is the difference of density between particle and fluid, and $\eta$ is the fluid viscosity. For a bacterial cell in water, $a$=1um, $g$=9.8m/s$^2$, $\Delta\rho$=80 kg/m$^3$, and $\eta$=10$^{-3}$ Ns/m$^2$. This yields to $V_s$=0.18 um/s.

At each time period $dt$, there are $n\delta V$ cells reaching the bottom surface, wit $n = N_0/(SH)$ the cell density, and $\delta V = V_S S\,dt$ the size of the micro-volume. This directly gives

$$N_s^g(t) = \frac{N_0}{H} V_S t$$

This is valid until the last cell reaches the surface (at time $t = H/V_S$). For $t > H/V_S$, $N_s^g(t)$ is constant and equal to $N_0$.

**b. Motile cells:**

Since the typical swimming speed (10 um/s) is much greater than the settling speed (Vs=0.18 um/s), we neglect the settling speed and only consider diffusion, with diffusion coefficient D, along the vertical axis z. Initially, cells are uniformly distributed between z=0 (bottom) and z=H (top). Cells get attached to the bottom surface at first contact. We estimate the number of cells sitting at the bottom surface over time using a first-passage approach.

Equation 91 of Meyer and collaborators[2] gives the first-passage probability density $Q$ of a particle starting at time t=0 at a position z and z=0 is the target position. Position z=H is reflective. The first-passage probability density is given with a renormalized time $\theta = t/\tau$, with $\tau = H^2/3D$ where H is the size of the box and D the diffusion coefficient.

$$Q(\theta, z) = \sum_{k=0}^{\infty} \frac{2\pi}{3}(k + \frac{1}{2})\sin[\frac{z}{H}(k + \frac{1}{2})\pi]\exp[-\frac{1}{3}(k + \frac{1}{2})^2\pi^2\theta]$$

If a particle is attached to the bottom surface at time $\theta^*$, it means it touched position $z = 0$ anytime between $\theta = 0$ and $\theta = \theta^*$. Starting from N particles at position $z$ at $\theta = 0$, only a fraction of these particles will have reached position $z = 0$ at time $\theta = \theta^*$. This fraction is $\alpha(\theta^*)N$, time-integration of the first-passage probability density $Q(\theta, z)$. Considering a uniform distribution of initial locations of the particles, we need to average this quantity across the interval $[0, H]$.

In the expression of $Q(\theta, x)$, time and space appear separately, which considerably facilitates the calculation.

Time-integration:

$$\int_0^{\theta^*} \exp(-\frac{1}{3}(k + \frac{1}{2})^2\pi^2\theta)d\theta = \frac{1}{\frac{1}{3}(k + \frac{1}{2})^2\pi^2}[1 - \exp(-\frac{1}{3}(k + \frac{1}{2})^2\pi^2\theta^*)]$$

Space-averaging:

$$\frac{1}{H}\int_0^H \sin(\frac{z}{H}(k + \frac{1}{2})\pi)dz = \frac{1}{H}\frac{H}{(k + \frac{1}{2})\pi}[1 - \cos((k + \frac{1}{2})\pi)] = \frac{1}{(k + \frac{1}{2})\pi}$$

Finally, it leads to:

$$\alpha(\theta) = \sum_{k=0}^{\infty} \frac{2}{(k + \frac{1}{2})^2\pi^2}\left[1 - \exp\left(-\frac{1}{3}(k + \frac{1}{2})^2\pi^2\theta\right)\right]$$

And by reverting to actual time:

$$\alpha(t) = \sum_{k=0}^{\infty} \frac{2}{(k+\frac{1}{2})^2\pi^2}\left[1 - \exp\left(-(k+\frac{1}{2})^2\pi^2 \frac{D}{H^2} t\right)\right]$$

And

$$N_s^d(t) = N_0 \sum_{k=0}^{\infty} \frac{2}{(k+\frac{1}{2})^2\pi^2}\left[1 - \exp\left(-(k+\frac{1}{2})^2\pi^2 \frac{D}{H^2} t\right)\right]$$

We can verify that all particles get attached to the bottom surface at long times:

$$\alpha(t=\infty) = \sum_{k=0}^{\infty} \frac{2}{(k+\frac{1}{2})^2\pi^2} = 1$$

The assumption of adhesion at first contact is strong and yields to over-estimating the number of cells attached at the bottom surface at a given time. We can explore its validity by calculating the average return time, the time needed for a cell to touch again the surface after one contact. Since $Q(\theta, z=0) = 0$ (a cell at position $z = 0$ immediately returns to position $z = 0$), we need to introduce a cut-off distance $x_0$ that defines the position of a cell that is about to leave the surface. Typically, $x_0 = 1\ \mu m$.

The average return time calculated as:

$\langle t \rangle = \frac{H^2}{3D}\langle \theta \rangle = \frac{H^2}{3D}\int_0^{\infty} \theta Q(\theta, x_0) d\theta$

Since $x_0 \ll H$, the sin term in the expression of $Q$ simplifies and it yields after calculations:

$$\langle t \rangle = \frac{Hx_0}{D}$$

With $H = 1\ mm$ , $x_0 = 1\ \mu m$, and $D = 10\ \mu m^2/s$, the average return time is $\langle t \rangle = 100\ s$. For a thinner channel (for instance $H = 50\ \mu m$), the average time can be much shorter (5 s).

#### III. Biofilm development kinetic described as a logistic growth

In order to formalize the hypothesis of the inoculated population abundance on the biofilm development kinetic, we made the hypothesis that the biofilm under flow could be reasonably described using a logistic equation as follows :

$$S(t) = \frac{K}{1 + \left(\frac{K - S_0}{S_0}\right)\exp(-bt)}$$

The equation captures the exponential growth of the dividing population size, S(t) and the saturation imposed by environmental factors such as nutrient limitation, toxic metabolites buildup or steric constrains[3]. b is the growth rate, K, the environmental carrying capacity, i.e. the maximal size the population can be reached, and $S_0$, the size of the initial population. Fig. S1A shows a series of curves S(t) generated using same arbitrary values of K and b and different values of $S_0$, from 0.1 to 100, which illustrates how initial abundance decrease delays biofilm development.

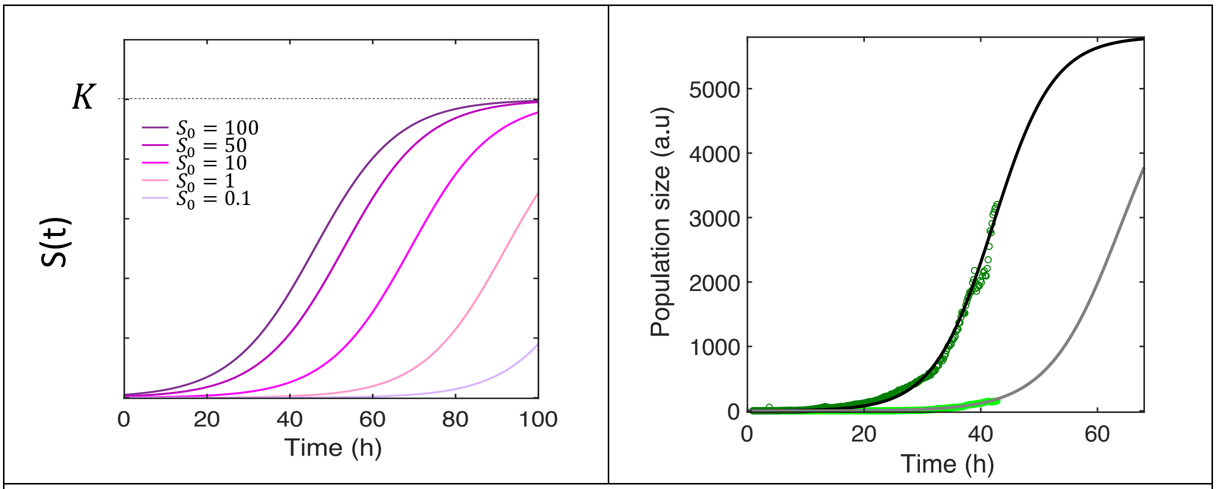


Figure **S3** (A) Ncurves (B) adjustments **Figure S1: Biofilm development phenomenologically fits logistic growth hypothesis**. (A) Typical logistic growth profiles within a three orders of magnitude range of initial population size $S_0$ (B) Adjustment of the experimental data with a logistic growth hypothesis; nonmotile cells biofilm data (dark green dot) are adjusted to a logistic profile with K= 5800 a.u.; b=0.19 $h^{-1}$(black line); motile cells data (light green dots) are adjusted using K=5800; b=0.17 $h^{-1}$.

To further investigate, the predictive quantitative power of our model, we tested the hypothesis of the logistic growth to account for our experimental data as a phenomenological guideline to score motile and nonmotile biofilms growth. Several parameter sets provided equally good quality adjustments with R-squared value >0.99. In particular, the parameter K accepted a whole range of values due the absence of description of the fully saturated level in the experiments. Nevertheless, assuming the same carrying capacity (5.8x$10^3$ in arbitrary units related to *E. coli* biofilm fluorescence signal) and similar growth rates (0.19 $h^{-1}$ and 0.17 $h^{-1}$ for nonmotile and motile biofilms, respectively, we obtained the adjustments displayed in Fig. S1B which provided an initial population sizes ratio (nonmotile to motile) of 12.

This ratio is approximately twice the ratio predicted by our sedimentation/diffusion model which suggests that the calculation slightly overestimates the number of cells that actually reach the surface by diffusion before flow starts. This might be due to the hypothesis of an adhesive first passage adhesion as explained above. Nevertheless, the whole picture strongly supports the hypothesis of development kinetic predominantly controlled by the initial abundance of the cells on the surface.

#### IV. Model predictions for geometric changes

We tested the predictions of our model as the characteristic height of the environment to be colonized changes. Diffusion coefficient and initial total population size remain constant. Fig. S4 shows that the relative abundance of the nonmotile to the motile cells on the surface increases over time up to a maximum, the time at which the maximum is reached also increasing with the characteristic height. It comes out that sedimentation versus diffusion discrepancy is essentially negligible for the small heights below 20$\mu$m, the relative abundance being very close to unity while it becomes significant in the millimeter range, reaching almost an order of magnitude and lasting longer. This behavior is expected to generate longer delays for the motile cells biofilm to develop as the height of the geometry increases. In the millimeter range, the difference between motile and nonmotile cells takes several tens of hours to completely relax.

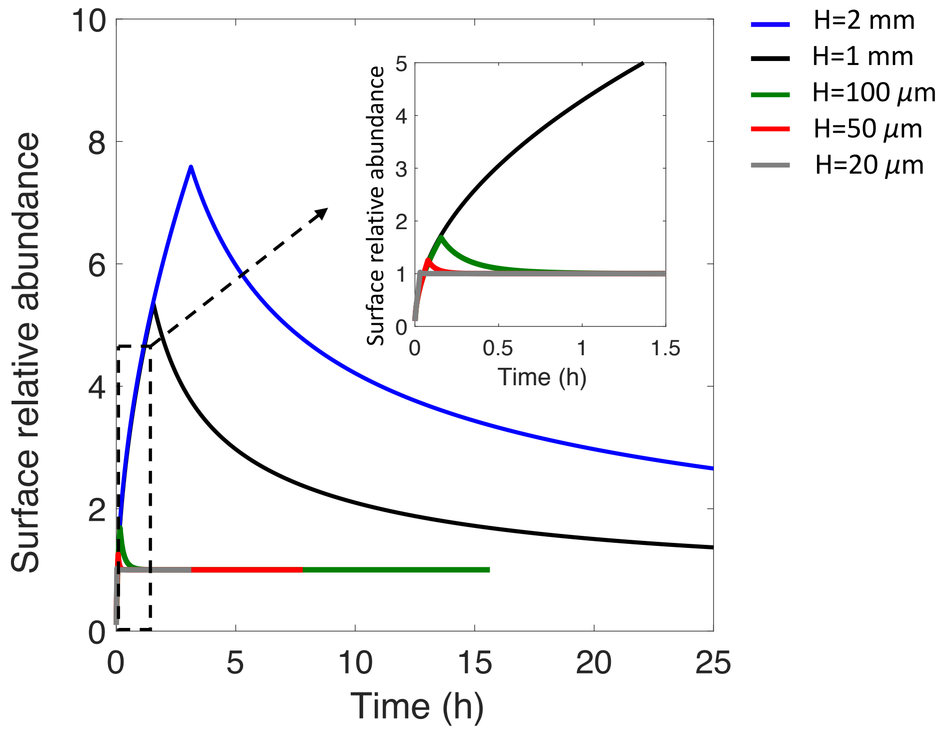


**Figure S4: Effect of the colonization geometry characteristic height.** The surface relative abundance of nonmotile to motile cells over time is shown for various height H of the environment to be colonized. ($D=10 \mu m^2/s$; $N_0=10^5$ cells).

#### V. Model predictions for diffusion coefficient changes

Bacterial swimming speed depends on several environmental factors and cell specific traits. In order to evaluate the impact of the flagellar motility efficiency on surface access, we examined the variation of the number of cells reaching the surface (first passage) (Fig. Sf(D)A) and the relative abundance compared to nonmotile cells (Fig. Sf(D)A) over time as diffusion coefficient varies at fixed device geometry (H=1000 $\mu$m) and total population size ($N=10^5$). It comes out that a higher diffusion coefficient accelerates the access to the surface. Therefore, in the limit of negligible settling of motile bacteria, the more motile bacteria will have a significant competitive advantage in surface colonization.

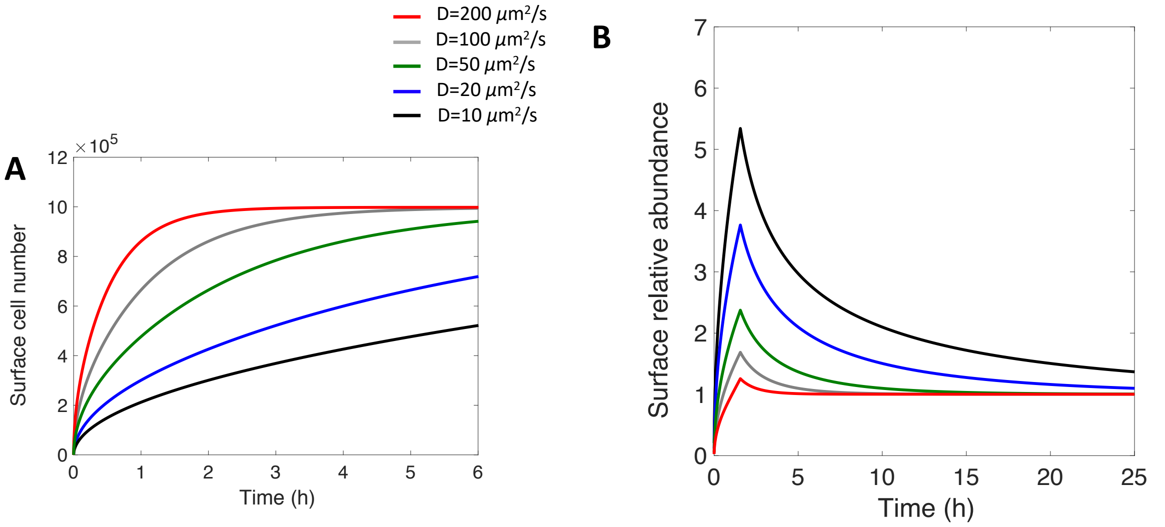


**Figure S5**: **Effect of diffusion coefficient on surface access kinetics.** (A) Number of cells reaching the surface over time for a range of diffusion coefficients from 10 to 200 μm$^2$/s (B) Corresponding relative surface abundance of nonmotile to motile cells. H=1mm, $N_0=10^5$ cells

#### VI. Colonization of the pre-established four-species biofilm

We also examined the impact of an already established community on motile and nonmotile *E. coli* ability to colonize the surface. To this purpose, we initiated the 4-species biofilm formation in the channel at time t=0 and performed *E. coli* injection after 8, 20 and 36 hours which corresponded to 4-species community first climax, beginning of the second growth phase and established dynamical equilibrium, respectively. We observed that *E. coli* installation under these conditions was negligible in any of these conditions, never overpassing the level of the fluorescence background produced by the pre-settled community. In each case, we measured FAST fluorescence 40 hours after *E. coli* injection in the pre-colonized channel and observed no difference between motile and nonmotile cell samples. We concluded that the initial surface coverage with a biofilm prevented *E. coli* installation regardless its motility.

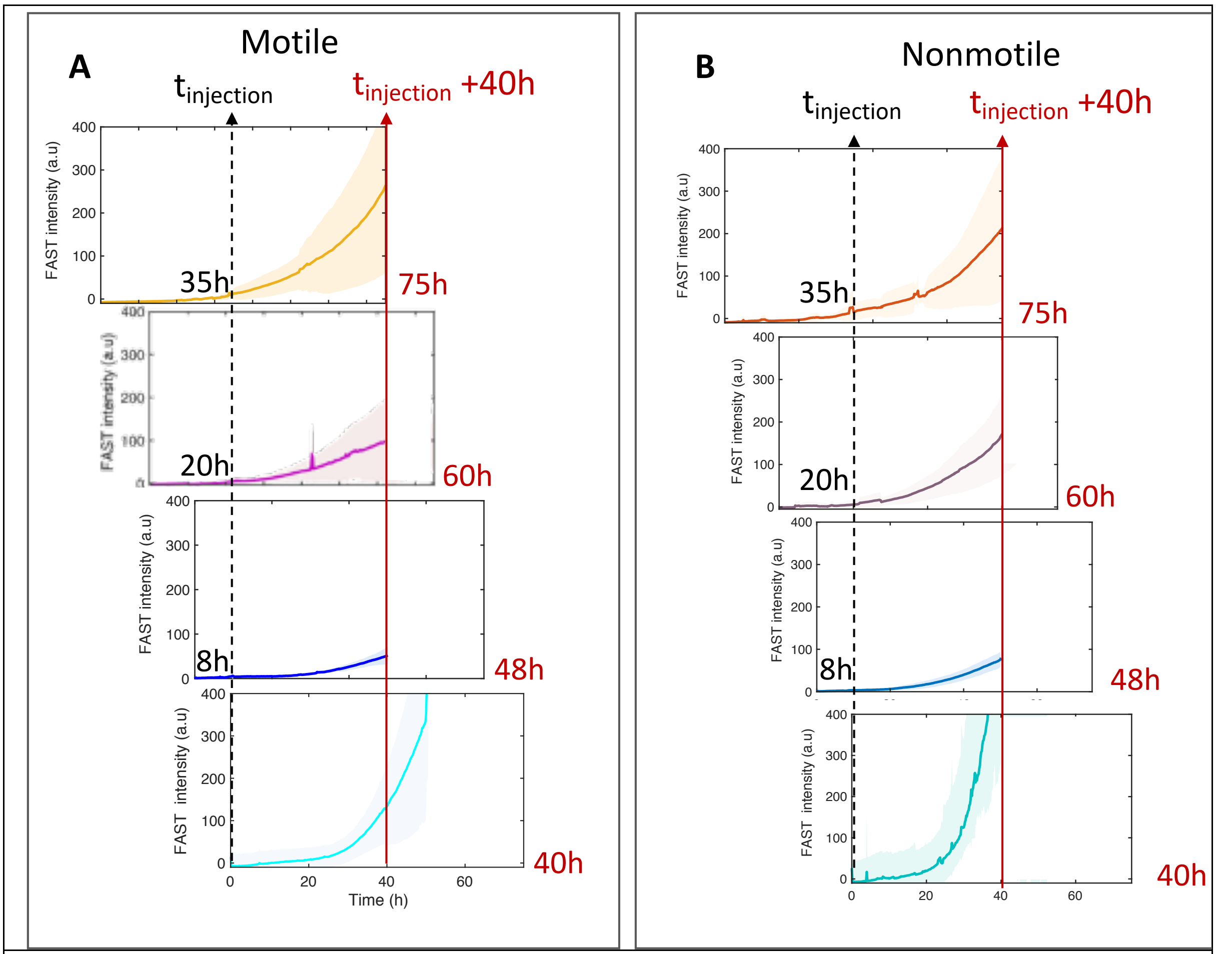


**Figure S6**: ***E. coli* fails to colonize surface with a pre-establish multispecies biofilm.** FAST fluorescence intensity on the channel surface — which report E. coli biofilm formation — is shown over the 40 hours of injection of *E. coli* in the channel where a 4S community has established for 0h, 8h, 20h and 35h from the bottom (0h) to the top (35h) of panel A for motile cells and panel B for nonmotile cells. The dashed vertical arrow indicates the injection time and the red arrow indicates time t=40h after injection. Only the co-colonizing situation corresponding to injection at time t=0 displays detectable *E. coli* colonization.

#### Supplementary data references


1 Barker, C. S., Pruss, B. M. & Matsumura, P. Increased motility of Escherichia coli by insertion sequence element integration into the regulatory region of the flhD operon. *J Bacteriol* **186**, 7529-7537, doi:10.1128/JB.186.22.7529-7537.2004 (2004).

2 Meyer, B., Chevalier, C., Voituriez, R. & Benichou, O. Universality classes of first-passage-time distribution in confined media. *Phys Rev E Stat Nonlin Soft Matter Phys* **83**, 051116, doi:10.1103/PhysRevE.83.051116 (2011).

3 Allen, R. J. & Waclaw, B. Bacterial growth: a statistical physicist's guide. *Rep Prog Phys* **82**, 016601, doi:10.1088/1361-6633/aae546 (2019).
